# Revisiting microbe-metabolite interactions: doing better than random

**DOI:** 10.1101/2019.12.10.871905

**Authors:** James T. Morton, Daniel McDonald, Alexander A. Aksenov, Louis Felix Nothias, James R. Foulds, Robert A. Quinn, Michelle H. Badri, Tami L. Swenson, Marc W. Van Goethem, Trent R. Northen, Yoshiki Vazquez-Baeza, Mingxun Wang, Nicholas A. Bokulich, Aaron Watters, Se Jin Song, Richard Bonneau, Pieter C. Dorrestein, Rob Knight

**Affiliations:** Department of Pediatrics, University of California, San Diego, La Jolla, CA, USA; Department of Computer Science & Engineering, University of California, San Diego, La Jolla, CA, USA; Center for Microbiome Innovation, University of California, San Diego, La Jolla, CA, USA; Collaborative Mass Spectrometry Innovation Center, University of California San Diego, La Jolla, CA, USA; Skaggs School of Pharmacy and Pharmaceutical Sciences, University of California San Diego, La Jolla, CA, USA; Department of Information Systems, University of Maryland Baltimore County, Baltimore, MD, USA; Department of Biochemistry and Molecular Biology, Michigan State University, East Lansing, MI, USA; Department of Biology, New York University, New York, 10012 NY, USA; Environmental Genomics and Systems Biology Division, Lawrence Berkeley National Laboratory, 1 Cyclotron Rd, Berkeley, CA, 94720, USA; DOE Joint Genome Institute, 2800 Mitchell Dr., Walnut Creek, CA, 94598, USA; Jacobs School of Engineering, University of California San Diego, La Jolla, CA, USA; The Pathogen and Microbiome Institute, Northern Arizona University, Flagstaff, AZ, USA; Department of Biological Sciences, Northern Arizona University, Flagstaff, AZ, USA; Flatiron Institute, Simons Foundation, New York, 10010 NY, USA; Computer Science Department, Courant Institute, New York, 10012 NY, USA; Center For Data Science, NYU, New York, NY 10008, USA; Department of Bioengineering University of California, San Diego, La Jolla, CA, USA

## Abstract

Recently, Quinn and Erb et al [1] made the case that when used correctly, correlation and proportionality can outperform MMvec when identifying microbe-metabolite interactions. We revisit this comparison and show that the proposed correlation and proportionality are outperformed by MMvec on real data due to their inability to deal with sparsity commonly observed in microbiome and metabolome datasets.

## II. RESPONSE

As shown in the original MMvec paper [2] and Quinn and Erb et al [1], scale invariance is key for recovering sensible microbe-metabolite interactions. However, contrary to the scale invariance argument made in the preprint [1], MMvec does not normalize the joint distribution *P*(***u*_*i*_**, ***υ*_*i*_**) between microbes and metabolites (microbe abundances are represented by ***u*_*i*_** and metabolite abundances are given by ***υ*_*i*_** for sample *i*). Instead, the MMvec algorithm attempts to model *P*(***υ*_*i*_**|***u*_*i*_**) with an inverse alr transform, a known compositionally coherent transform that satisfies scale invariance [3]. This approach is more similar to a conditional version of approach B in the preprint rather than approach A [1]. Because of this, our method does not have the stated problem that microbe and metabolite abundances compete for probability mass in the normalized distribution: “the abundance of microbe 1 is limited by the abundance of microbes 2-to-M, but is in no way limited by the abundance of metabolites 1-to-N”.

We relied on simulated data in [2] for the purpose of argument, and to illustrate some of the principles in the paper clearly and without the distractions typically present in real data. However, simulated data will always have limitations because of the inability to model unknown features of the real system, or because of deliberate simplifications that clarify key points in the model system. Therefore, a crucial aspect of the MMvec manuscript was to test performance both on simulations and on real data. Reuse and re-purposing requires a thorough understanding of the simulated data. Performance on real data is the ultimate test of methods, and any simulated data experiment should always be accompanied with evaluation using real data, which was not done in the commentary. Accordingly, we applied the same proportionality-based scripts described in the preprint [1], and evaluated them on one of the real datasets we used in the MMvec paper.

A major obstacle to analyzing real-world microbiome and metabolomics data is sparsity. Traditional compositional methods such as the proposed clr transform cannot automatically deal with zeros, and require imputation as a preprocessing step. This imputation adds bias, and is impractical for the sparse datasets typically encountered [4, 5]. Microbiome and untargeted metabolomics datasets are generally sparse; in large studies, such as the American Gut Project [6], the sparsity for stool samples alone is 99.946%. MMvec was designed to handle very sparse data using bootstrapping and a multinomial likelihood function, without any imputation. With the desert biocrust soils dataset (sparsity of 51%; [7]) that was used in the MMvec publication, we observe that MMvec outperforms the newly proposed linear methods dramatically (Fig. 1).

**FIG. 1.**
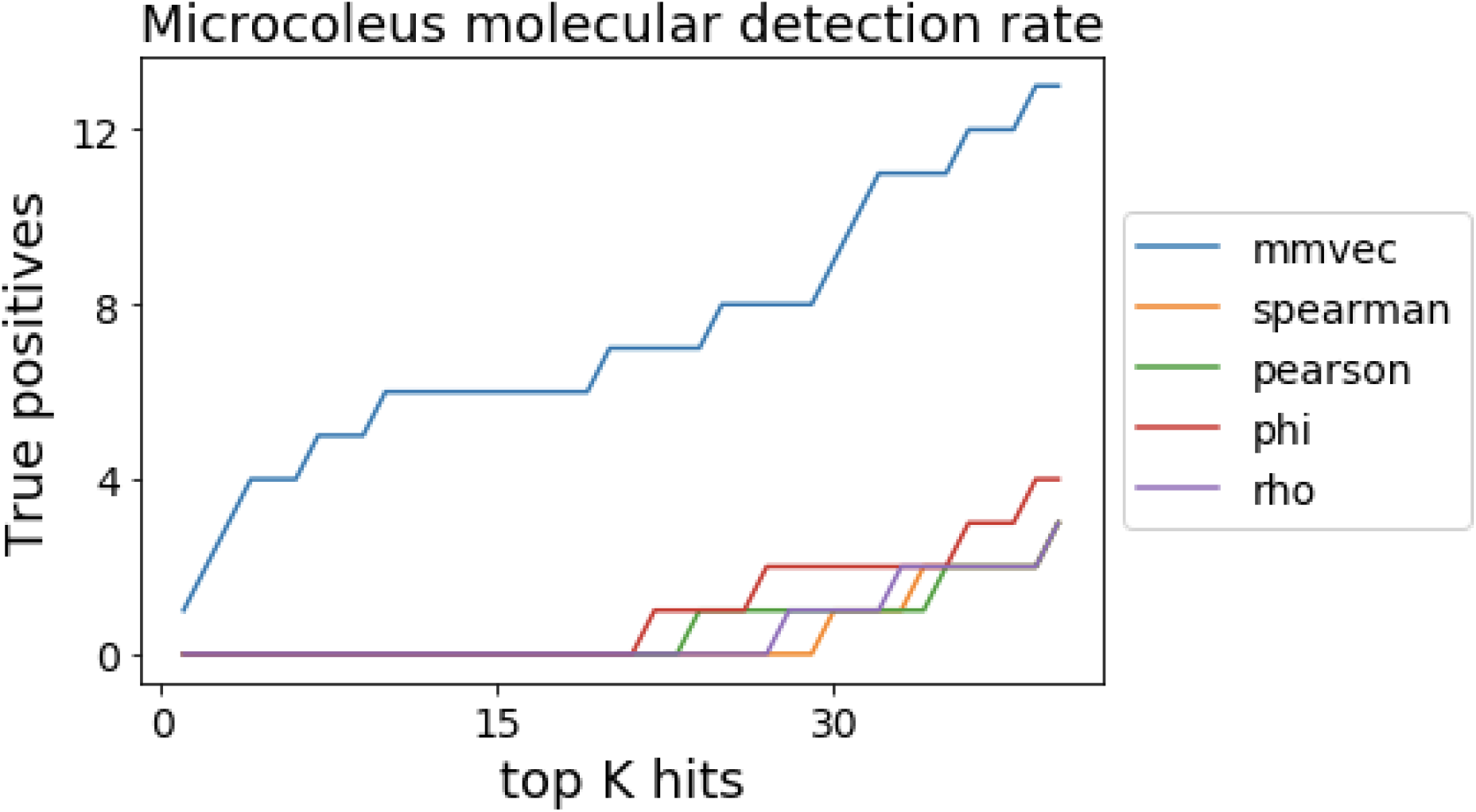
A comparison of mmvec to metrics proposed by Quinn and Erb [1]. These proposed metrics include Spearman, Pearson, phi and rho applied after a clr transformation [3].

Our findings here are consistent with the key conclusions in a separate preprint [8]. Specifically, they showed that their neural network approach also outperformed linear methods. Coincidentally, their paper evaluated an IBD data set [9] similar to one of the examples in the MMvec paper, and were largely concordant with our MMvec findings. In fact, the abstract states that: “In this paper, we propose a sparse neural encoder-decoder network to predict metabolite abundances from microbe abundances. Using paired data from a cohort of inflammatory bowel disease (IBD) patients, we show that our neural encoder-decoder model outperforms linear univariate and multivariate methods in terms of accuracy, sparsity, and stability.” Although their proposed underlying model achieves a different goal, our MMvec findings are consistent with those statements.

Contrary to the argument in the preprint [1] regarding the complexity of neural networks, the MMvec model [2] is not much more complex than the proposed regression techniques; it is a simple one-layer neural network without complex activation functions, which is in effect a two-stage log-bilinear regression. MMvec has even less computational complexity than the proposed proportionality metrics. Proportionality requires the estimation of *O*(*NM*) parameters, one parameter for each microbe-metabolite interaction, which can easily result in the need to estimate millions of parameters for systems with only thousands of microbes and thousands of metabolites. This large number of parameters is problematic, but was not discussed in the commentary [1]. In contrast, MMvec assumes a low-dimensional latent representation requiring the estimation of *O*(*Nk* + *Mk*) parameters. In practice, this amounts to estimating only thousands of parameters if the latent dimensionality is small (i.e. *k* < 10).

Methods similar to MMvec have been successful at the task of learning word embeddings. Since Mikolov et al. [10], these models have been designed with an emphasis on practical methods for learning useful embeddings at scale, rather than on perfectly modeling the data distribution. Imperfect modeling assumptions (if they do indeed exist in our case) do not prevent a method’s successful use for a particular task. The quote from George Box comes to mind, “all models are wrong. Some models are useful”.

MMvec is only one tool in the arsenal of correlative methods. It is not perfect for every correlation type or dataset, and is not a one size fits all “magical” solution [11]. However, we have found that MMvec is a powerful discovery tool, as demonstrated by the other real datasets we evaluated in the original article, and by its wide use; the tool has already been downloaded >1200 times, and several papers based on it have already been submitted [12, 13], with many more in the works. It is critical that we provide accurate guidance to the community so that scenarios where one method works better than others are well understood. Fundamentally, the argument that simple linear methods outperform neural networks was not supported in the commentary, because only dense simulation datasets were evaluated. We appreciate the communication on the topic to the extent that it helps the community better understand the advantages of the different approaches, and prompts the community to continue to innovate in this area.

## III. SOFTWARE AVAILABILITY

The analysis can be found under : https://github.com/knightlab-analyses/multiomic-cooccurrences/blob/rebuttal/ipynb/Figure3-rerun.ipynb

